# TBPL2-dependent transcription controls RNA stability and cellular organization in growing oocytes

**DOI:** 10.64898/2026.09.18.752403

**Authors:** Emmanuel García Sánchez, Dylane Detilleux, Claire Richard, Louise Couturier, Céline Ziegler-Birling, Nadia Messaddeq, Fabienne Mauxion, Bertrand Séraphin, László Tora, Stéphane D. Vincent

## Abstract

During oocyte growth, the marked increase in cell size is accompanied by robust RNA polymerase II (Pol II) transcription, generating transcripts that are either translated into proteins or stored as part of the maternal transcriptome. While the RNA decay machinery is known to critically regulate the maternal transcriptome during oocyte maturation, its contribution during the growth phase remains poorly understood. We previously identified an oocyte-specific transcription machinery dependent on the TATA-binding protein paralog TBPL2, which is required for oocyte growth. To investigate how the absence of TBPL2-mediated transcription affects the cellular state of the oocyte, we characterized the phenotypic consequences of the *Tbpl2^-/-^* mutation in growing oocytes. Our data reveal that both nuclear and cytoplasmic organizations are profoundly disrupted in the absence of TBPL2-mediated transcription. Given that cytoplasmic organization is closely linked to RNA storage capacity, we examined the expression and localization of proteins involved in mRNA regulation, stability and storage. We observed a marked impairment in the redistribution of these key factors in *Tbpl2^-/-^* oocytes. To assess the impact of TBPL2 on RNA storage, we analyzed poly(A) tail length and found a global increase in polyadenylation in mutant oocytes. Gene-specific poly(A) tail analyses revealed differential mRNA changes, with shortened tails for a downregulated transcript and elongated tails for an upregulated one, suggesting stabilization of the latter. Collectively, our findings indicate that TBPL2-mediated transcription is essential for maintaining proper cellular organization in growing oocytes, at least in part by ensuring the functional integrity of the RNA decay machinery.

## Introduction

In female mice, oogenesis is initiated during embryonic development, with the proliferation of oogonia and their entry in meiosis I around embryonic day (E) 13.5. Remarkably, all the primary oocytes are arrested at the end of prophase I (dictyate stage) and will remain arrested throughout adulthood unless they are selected for ovulation. These primary oocytes are initially interconnected, forming germ cysts (for a review, see (Pepling, 2006)). At birth, the breaking of germ cysts leads to the formation of primordial follicles, each composed of a single arrested primary oocyte associated with few somatic follicular cells. These primordial follicles constitute the reserve of germ cells that will support the reproductive life of the females. Regularly, a subset of these primordial follicles is primed for further differentiation, a process called folliculogenesis.

Folliculogenesis is the process by which follicles increase in size and complexity, involving the proliferation of follicular cells, and progress from a primordial stage to a pre-ovulatory stage (for a review, see (Tora & Vincent, 2021)). The primary oocyte, still arrested at the dictyate stage, also undergoes changes during folliculogenesis, which is divided in two phases. The first phase occurs from the primary to the pre-antral follicular stage and is associated with the oocyte growth. By the end of the pre-antral follicular stage, the oocyte growth is complete and the oocyte enters the second phase which involves oocyte maturation. During this second phase, maturation leads to the resumption of meiosis culminating in ovulation. In mammals, there are two waves of folliculogenesis. The first wave occurs after birth in the medullar region of the ovary in mice, while the second wave occurs in the cortical region of the ovary and will support the whole reproductive life of the females (Mork *et al*, 2012; Zheng *et al*, 2013).

The intense growing phase is associated with an increase in transcription activity in the growing oocytes, allowing the accumulation of transcripts that are stored and constitute the maternal transcriptome. As transcription stops at the end of the growing phase and will only resume after fertilization, post-transcriptional regulation of the maternal transcriptome is crucial to support maturation, ovulation and initiation of development. A key mechanism to control the storage and the translatability of the transcripts is the regulation of the poly(A) tail length (Bachvarova, 1992; Paynton & Bachvarova, 1994). During oocyte growth, several compartments are established, associated with the storage of maternal proteins and/or mRNAs. First, the formation of highly ordered lattice-like structures called cytoplasmic lattices (CPLs) has been observed in the cytoplasm of the growing oocyte (Burkholder *et al*, 1971) (for a review, (Giaccari *et al*, 2024)). These CPLs are a matrix composed of proteins and RNA and play a major role in the storage of ribosomes (Yurttas *et al*, 2008) and of maternal proteins (Jentoft *et al*, 2023). More recently, the CPL structure was resolved showing that it also stores tubulin heterodimers and an array of ubiquitination proteins (Kılıç *et al*, 2026; Li *et al*, 2026; Liu *et al*, 2026; Chi *et al*, 2026). Interestingly, these structures are also involved in the storage of maternal mRNA via their interaction with the non-specific RNA binding protein YBX2 (MSY2), which is involved in the stabilization of maternal RNA in oocytes (Liu *et al*, 2017). This compartment is associated with a second compartment in the cortical region of the oocytes, called the subcortical maternal complex (SCMC) (Li *et al*, 2008) which is important for the regulation of the actin dynamics (Yu *et al*, 2014). These two structures share many maternal proteins such as PADI6, TLE6, MATER/NLRP5, FILIA/KHDC3 and OOEP (for a review see (Bebbere *et al*, 2021)) and are now considered as a single structure which organizes the cytoplasm of the growing oocyte (Jentoft *et al*, 2023; Chi *et al*, 2026; Kılıç *et al*, 2026; Li *et al*, 2026; Liu *et al*, 2026). In the same subcortical region, subcortical aggregates (SCA) have been described to be associated with RNA as a result of the dissolution of the P-bodies during oocyte growth (Flemr *et al*, 2010). Recently, another compartment associated with mitochondria and the storage of RNA, established during oocyte growth, has been described, called the mitochondria associated ribonucleic domain (MARDO) (Cheng *et al*, 2022).

Interestingly, the intense transcription activity during oocyte growth is associated with a transition in the RNA polymerase II (Pol II) transcription initiation machinery. The initiation of Pol II transcription starts with the nucleation of the Pre-Initiation Complex (PIC) on the core promoters. A functional PIC is composed of the Pol II and 6 General Transcription Factors (GTF) and among these GTF, TFIID, composed of the TATA binding protein (TBP) and 13 TBP-associated factors (TAFs), is the first complex to recognize and to bind to the core promoter sequences (for a review, see (Malik & Roeder, 2023)). At the beginning of oocyte growth, the closest homolog of TBP in vertebrates, TBPL2/TBP2/TRF3 (hereafter called TBPL2) starts to be expressed (Gazdag *et al*, 2007) and transcription initiation is mediated independently of a holo-TFIID by a complex composed of TBPL2 and the GTF TFIIA (Yu *et al*, 2020). In *Tbpl2^-/-^* females, oocyte growth is blocked between the primary and the secondary follicular stages, associated with a strong reduction in H3K4me3 and barely detectable serine 2 phosphorylation on C-terminal repeat (CTD) of Pol II largest subunit RPB1 (Gazdag *et al*, 2009). No increase in apoptosis could be detected by TUNEL in the mutant oocytes, but the follicles’ structure was disorganized at 6 weeks (Gazdag *et al*, 2009).

In this study, we investigated the cellular phenotype of *Tbpl2^-/-^*mutant growing oocytes to assess the impact of the lack of TBPL2-mediated Pol II transcription on oocyte development. Our data indicate that both the nuclear chromatin and cytoplasmic organization are profoundly disrupted in mutant oocytes. Given that cytoplasmic organization is closely linked to the capacity of the oocyte to store RNA, we further analyzed the expression and localization of proteins involved in mRNA regulation, stability, and storage, and observed impaired redistribution of these factors in mutant oocytes. We next investigated the consequences for RNA storage and, unexpectedly, detected a global increase in poly(A) signal in the absence of TBPL2. Gene-specific analyses revealed that poly(A) tail length is reduced in downregulated transcripts but increased in upregulated transcripts, strongly suggesting enhanced stabilization of upregulated transcripts in the absence of TBPL2. Together, these findings demonstrate that both RNA metabolism and cellular organization are severely perturbed in *Tbpl2^-/-^* mutant growing oocytes, establishing a TBPL2-dependent coordination of transcription-RNA stability coupling.

## Results

### Tbpl2^-/-^ oocytes have an abnormal structure

The disorganization of the *Tbpl2^-/-^* mutant follicle was not yet precisely evaluated in earlier studies (Gazdag *et al*, 2009). To get more insight into the defects in the *Tbpl2^-/-^* mutant growing oocytes, we carried out histological analyses at different times: post-natal day (PN) 14, 6 weeks, 8 weeks and 32 weeks (Figure 1). At PN14, the *Tbpl2^-/-^* mutant ovaries are much smaller, due to the reduced development of the mutant follicles associated with the accumulation of primary follicles and failure to develop secondary follicles (Figure 1A). While the size of the WT ovaries increased with age, the mutant ovaries remained smaller at all stages analyzed (Figure 1B-D). The depletion of follicles in the mutant ovaries, already reported at 6 weeks (Gazdag *et al*, 2009) (Figure 1B), became very severe at 8 and 32 weeks, with only few follicles with an abnormal structure (Figure 1C,D, arrow). The residual follicles remaining in the mutant at 8 weeks were characterized by a loss of the interaction between follicular cells and the oocyte (Figure 1E). Analysis at 32 weeks showed that some oocytes were still present in the mutant ovaries, but the remaining follicles are completely disassembled and the oocytes lose their contact with the follicular cells (Figure 1E). Interestingly, we observed that in the young ovaries, the cortical folliculogenesis was impaired (Figure 1F) and that oocytes persisted in the mutant ovaries but started to lose their connections with the follicular cells (Figure 1G). In addition to their abnormal structure, the nuclei of the mutant oocytes were significantly bigger compared to the control counterpart at all ages (Figure 1H) associated with a decrease in the cytoplasmic diameter and an increase of the nuclear/cytoplasm ratio (Figure S1A,B). Altogether, these data indicate that in addition to the impairment of the primary to secondary follicles transition, absence of TBPL2-mediated transcription impacts the structure of the follicles and the oocyte cellular organization.

**Figure 1:**
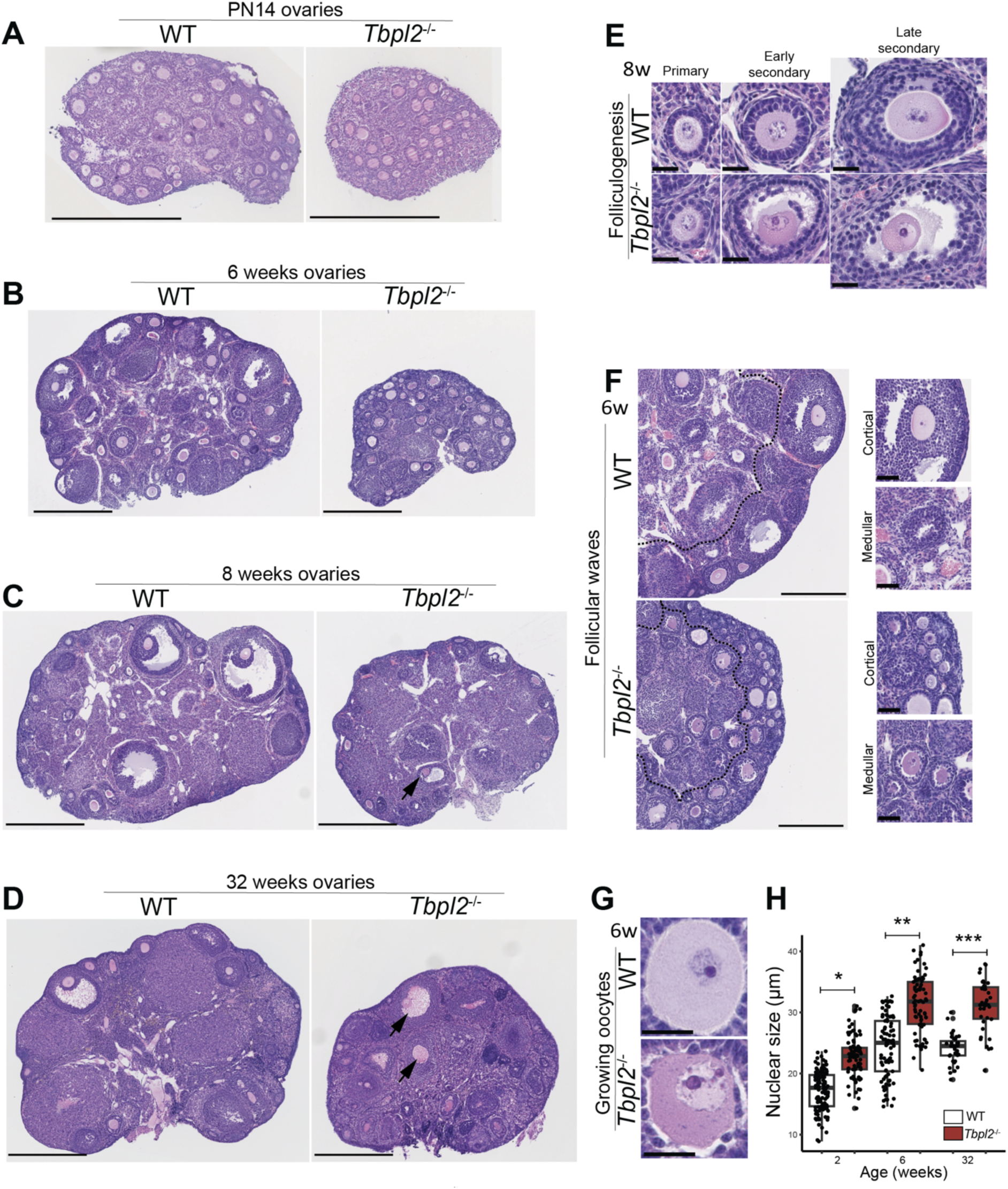
Histological analysis of *Tbpl2*^-/-^ ovaries. **(A-D)** Comparative hematoxylin-eosin (HE) study of WT and *Tbpl2*^-/-^ovaries at post-natal (PN) 14 days (A), 6 weeks (B), 8 weeks (C) and 32 weeks (D) old females. Scale bars: 500 µm, arrows in (C,D) point to abnormal follicles. (**E**) Folliculogenesis at primary, early secondary and late secondary follicular stages from WT and *Tbpl2*^-/-^ ovaries at 8 weeks. Scale bars: 25µm. (**F**) Follicular waves at 6 weeks in WT and *Tbpl2*^-/-^ mutant ovaries. Dashed line: delimitation of the cortical and medullar follicular waves. Magnification of cortical versus medullar follicles. Scale bars: 250 µm and 50 µm (magnifications). (**G**) Abnormal structure of the *Tbpl2*^-/-^ oocytes at 6 weeks. Scale: 25 µm. (**H**) Quantification of the size of the oocytes nuclei at PN14, 6 and 32 weeks. Nuclei of *Tbpl2^-/-^*oocytes are significantly bigger compared to WT counterparts. Wilcoxon test, *: *p*-value ≤ 0.05, **: *p*-value ≤ 0.01, ***: *p*-value ≤ 0.001.

### Abnormal chromatin organization in Tbpl2^-/-^ mutant oocytes

As the larger size of the mutant nuclei could be the result of a failure to properly pack the chromatin in the *Tbpl2^-/-^* mutant oocytes, we evaluated the presence and levels of markers associated with chromatin structure. Previous study has indicated that H3K4me3 histone marks are decreased in the mutant oocytes at 6 weeks (Gazdag *et al*, 2009). In order to investigate the potential defects at an early step, we carried out immunolocalization in isolated PN14 oocytes. We first analyzed histone marks associated with compacted chromatin (facultative heterochromatin; H3K27me3 and constitutive chromatin; H3K9me3) (for a review, see (McCarthy *et al*, 2023)) (Figure 2). While only a moderate decrease could be observed in the H3K27me3 expression pattern (Figure 2A,B), the H3K9me3 signal was completely absent in mutant oocytes (Figure 2C,D), strongly suggesting that the chromatin is globally decompacted in the absence of TBPL2. Expression of the HP1beta protein associated with chromatin compaction (for a review, see (McCarthy *et al*, 2023)) did not appear obviously perturbed in the mutant nuclei as foci were still present, however, there was a significant diffuse signal in the nucleus associated with an increase in the global intensity of HP1β, suggesting that the chromatin was not comparably compacted between the control and the mutant oocytes (Figure 2E,F). Finally, analysis of 5-methyl cytosine (5mC) using anti 5-methyl cytosine antibody showed a severe decrease intensity (Figure 2G,H), in agreement with the fact that this mark is deposited co-transcriptionally (Veselovska *et al*, 2015) and that TBPL2 is required for active transcription in the oocytes (Yu *et al*, 2020). Altogether these data indicate that there is a correlation between the decompacted state of the chromatin and the increased size of the nuclei in the *Tbpl2^-/-^*mutant oocytes.

**Figure 2:**
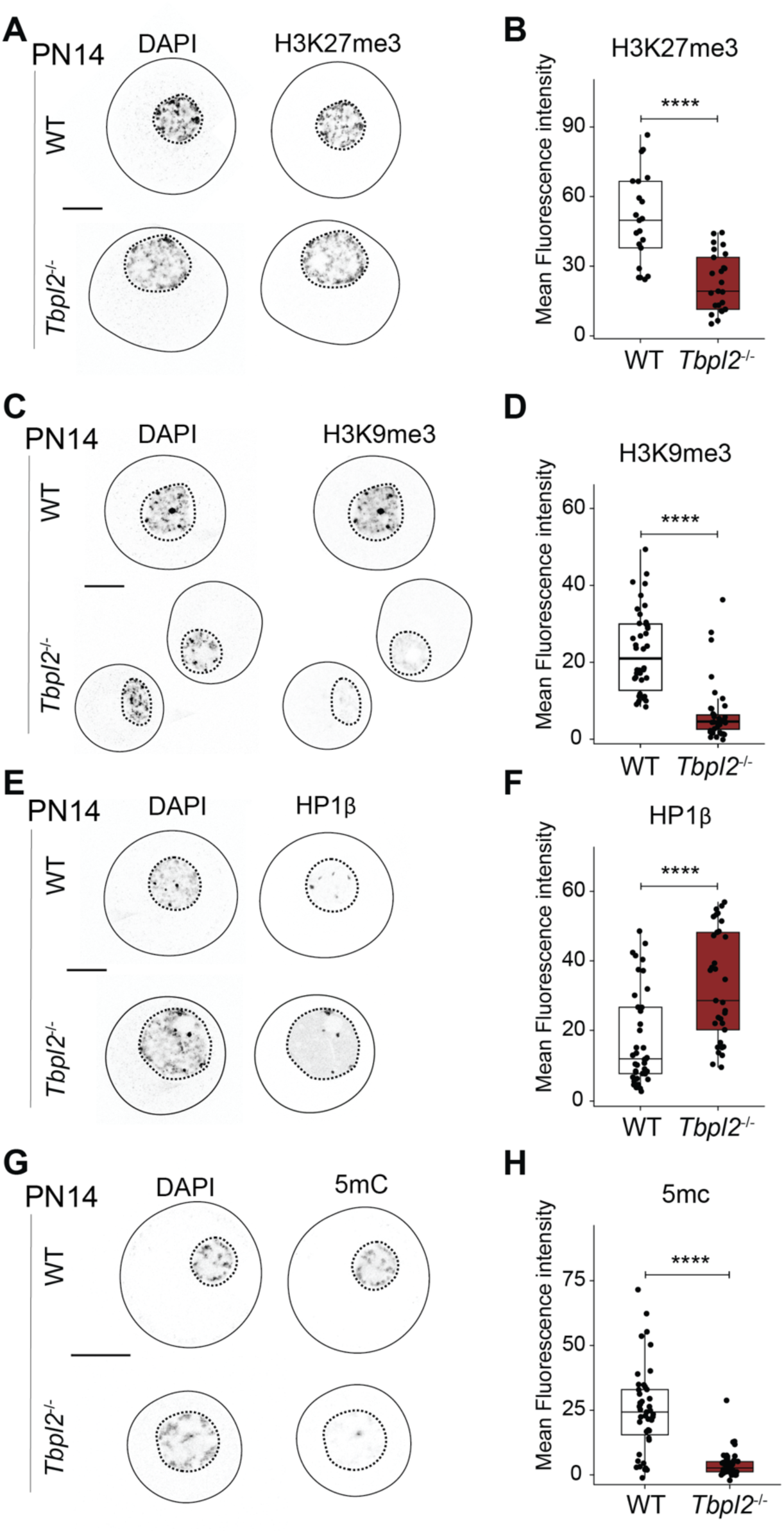
Abnormal chromatin organization in *Tbpl2*^-/-^ oocytes. **(A-G)** Negative grayscale pictures of immunofluorescence (A,C,E,G) and mean fluorescence intensity quantification of the nuclear signal (B,D,F,H) on growing WT and *Tbpl2*^-/-^ oocytes at PN14 using antibodies against: facultative heterochromatin histone mark H3K27me3 (A,B, N=2, n_WT_=21, n_Tbpl2-/-_=24), constitutive chromatin histone mark H3K9me3 (C,D, N=3, n_WT_=42, n_Tbpl2-/-_=40), heterochromatin-associated factor HP1β (E,F, N=3, n_WT_=39, n_Tbpl2-/-_=33) and 5mC modification (G,H, N=4, n_WT_=45, n_Tbpl2-/-_=37). Pictures displayed are single planes, scale bars: 20 µm, Wilcoxon test, *: *p*-value ≤ 0.05, **: *p*-value ≤ 0.01, ***: *p*-value ≤ 0.001, ****: *p*-value ≤ 0.0001.

### Cytoplasmic organization is severely perturbed in the Tbpl2^-/-^ mutant oocytes

Next, we analyzed the expression and localization of different cytoskeletal proteins, including TUBULIN, F-ACTIN and DESMIN, in oocytes at PN7 (primary follicles, initiation of growth) and PN14 (secondary follicles, growing oocytes) stages. At PN7, expression levels and localization of β-TUBULIN (Figure 3A) and F-ACTIN (Figure 3B) were comparable between controls and *Tbpl2^-/-^* mutant oocytes with their apparent accumulation in the subcortical region. In the PN7 controls, expression of the intermediate filament protein DESMIN was localized in a cluster close to the nucleus and this pattern was similar in the absence of TBPL2 (Figure 3C). Altogether, these data indicate that at the early stage of oocyte growth, the cytoskeleton organization is not affected in the *Tbpl2^-/-^* mutant oocytes. Remarkably, absence of TBPL2 expression severely affects the re-organization of the cytoskeleton during oocyte growth as observed at PN14. While both TUBULIN and ACTIN are strongly enriched in the subcortical domain in the WT growing oocytes at PN14, in the *Tbpl2^-/-^*mutant oocytes, no such subcortical localization is achieved (Figure 3D-F). More precisely, these proteins remained clustered in close contact with the nucleus (Figure 3D,F). During growth in WT oocytes, the initial clustered DESMIN domain observed at PN7 is reorganized in a more diffuse pattern of cytoplasmic aggregates at PN14 (Figure 3G). This drastic reorganization is not observed in the *Tbpl2^-/-^*mutant oocytes as DESMIN containing cytoplasmic aggregates are still clustered in close apposition to the nucleus (Figure 3G,H). As the cytoskeleton was initially similar in both WT and the *Tbpl2^-/-^*mutant oocytes at the initiation of growth, these data indicate that the defects observed were a consequence of the lack of TBPL2 and also suggest that the cytoplasm cytoskeleton does not reorganize in the *Tbpl2^-/-^* mutant oocytes. To get more insight into the re-organization we observed during the growth phase, we investigated two organelles, the Golgi and the mitochondria. The analysis of the Golgi marker GM130 showed that the Golgi is following a similar reorganization as DESMIN, with an initial accumulation in a cluster close to the nucleus at PN5 (before initiation of the growth) and at PN7 (Figure S2A, Figure 4A) in both WT and mutant oocytes, and became more diffuse with the appearance of aggregates enriched around the nucleus but also present in the whole cytoplasm at PN14 in WT oocytes (Figure 4B). As observed for DESMIN, the Golgi fails to be reorganized and is limited to cytoplasmic aggregates clustered in the center of the oocytes, in close apposition of the nucleus in the *Tbpl2^-/-^* mutant oocytes (Figure 4A,B). Then we analyzed the distribution of the mitochondria during oocyte growth. It has been shown that during oocyte growth, the mitochondria are initially present as cytoplasmic clusters in a homogenous pattern (Cheng *et al*, 2022), as we observed in the control growing oocytes at PN14 (Figure 4C, Figure S2B). In contrast, in the absence of TBPL2, the distribution of the mitochondria was not homogenous as they were clustered in close apposition to the nucleus (Figure 4C). Interestingly, we observed that in the mutant oocytes, the Golgi and the mitochondria were regrouped together compared to the WT oocytes (Figure S2B). The association of a more centrally located Golgi surrounded by mitochondria, in association with one side of the nucleus is strikingly reminiscent of the structure of the Balbiani body commonly found in the oocytes of many organisms, including zebrafish or human, but surprisingly not in the mouse (Jamieson-Lucy & Mullins, 2019; Dhandapani *et al*, 2024).

**Figure 3:**
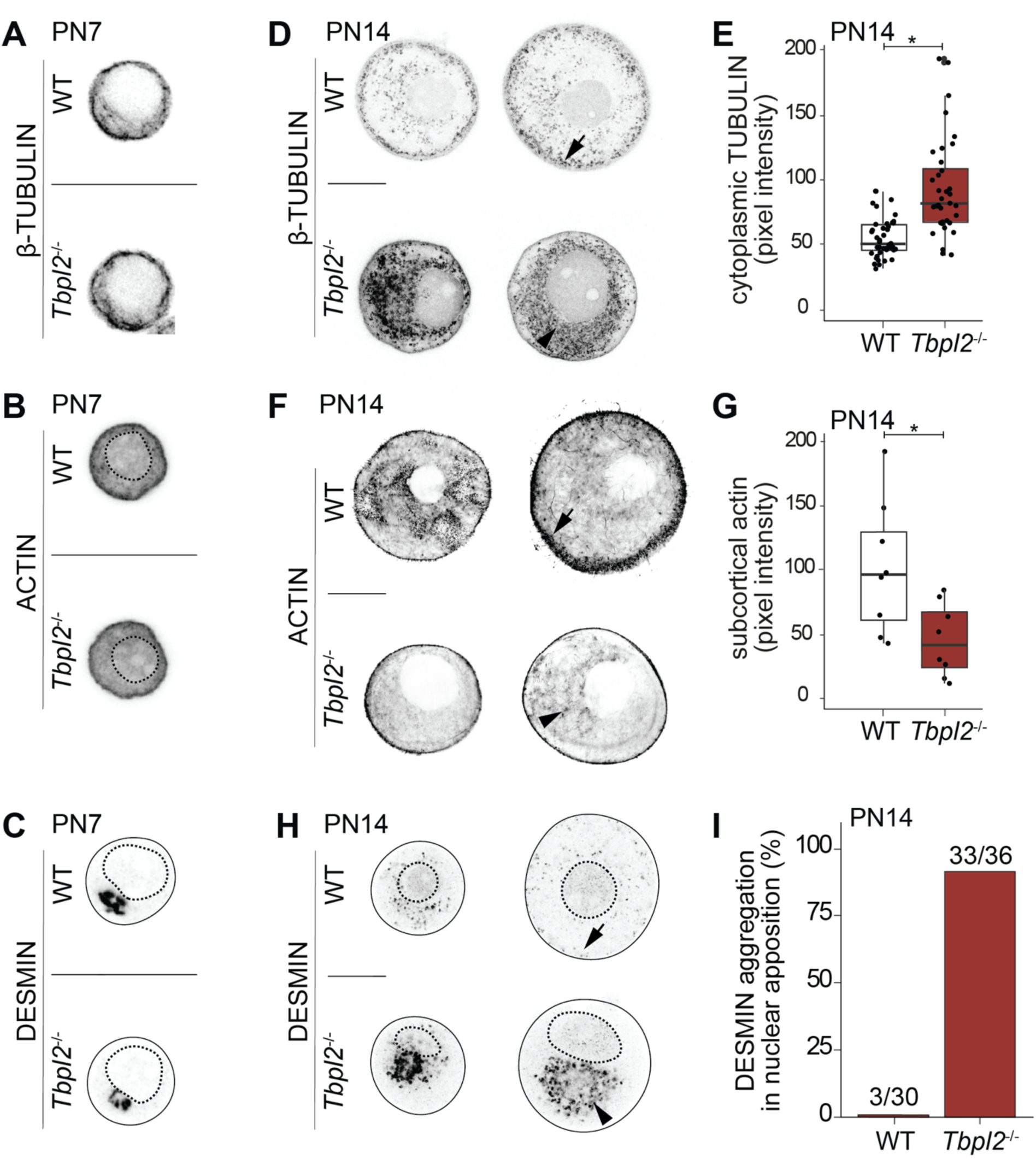
Disorganization of the cytoskeleton in *Tbpl2*^-/-^ mutant oocytes. **(A-C)** Detection of β-TUBULIN (A), ACTIN (B) and DESMIN (C) in growing WT and *Tbpl2*^-/-^ mutant oocytes at PN7. **(D-I)** Detection of β-TUBULIN (D, quantification of cytoplasmic β-TUBULIN in E, n_WT_=42, n_Tbpl2-/-_=36), ACTIN (F, quantification of subcortical ACTIN in G, n_WT_=8, n_Tbpl2-/-_=8) and DESMIN (H, quantification of DESMIN aggregation in nuclear apposition in I, n_WT_=30, n_Tbpl2-/-_=36) in growing WT and *Tbpl2*^-/-^ mutant oocytes at PN14. Arrows indicate cortical signal and arrowheads show cytoplasmic aggregation in close apposition to the nucleus in (D,F,H). The negative grayscale pictures displayed are single planes, scale bars: 25 µm, Wilcoxon test, *: *p*-value ≤ 0.05.

**Figure 4:**
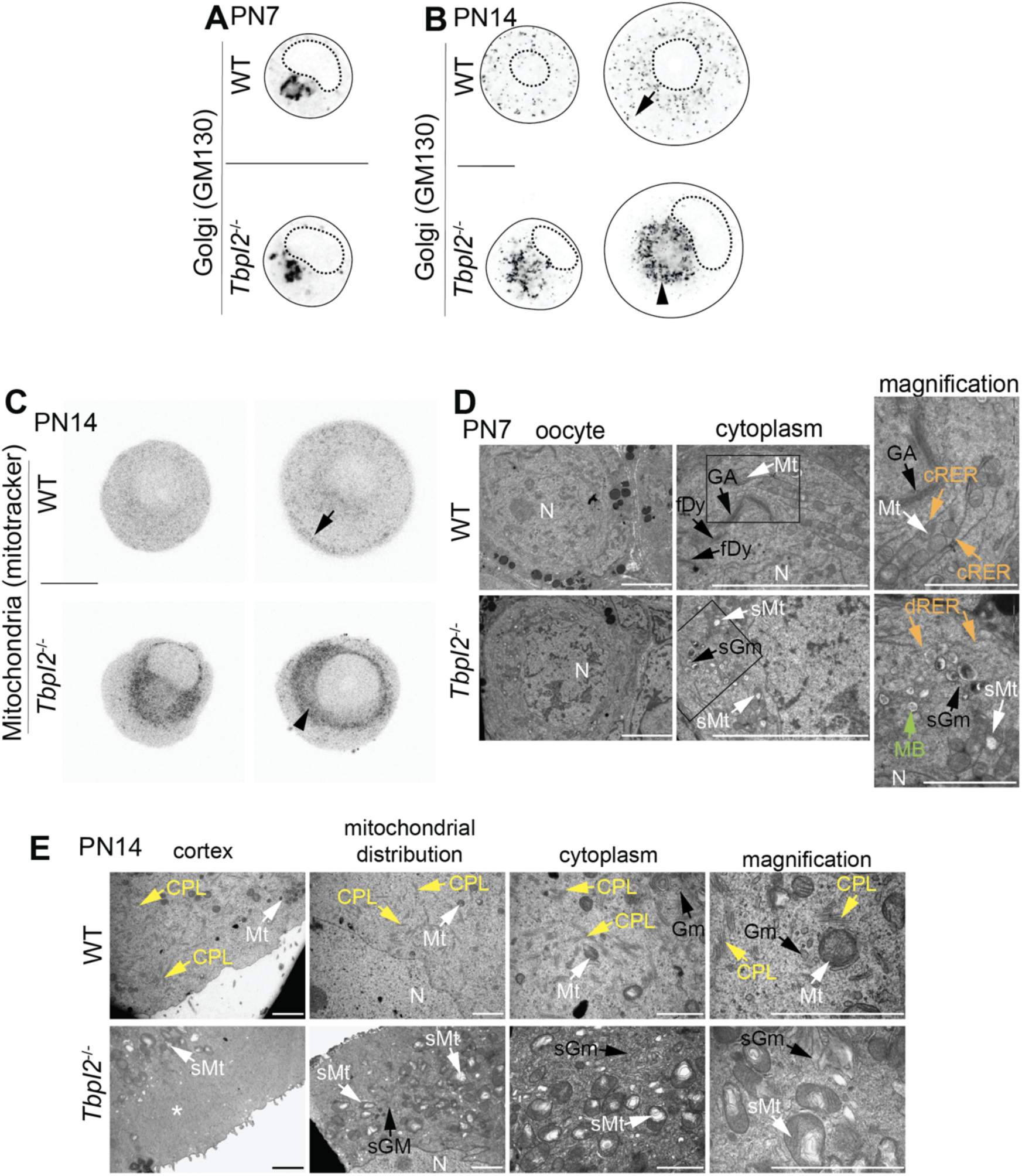
Abnormal cytoplasmic organization in *Tbpl2*^-/-^ mutant oocytes. (**A**,**B**) Negative grayscale pictures of immunofluorescence on growing WT and *Tbpl2*^-/-^oocytes: Golgi-associated protein GM130 at PN7 (A) and PN14 (B). (**C**) Mitochondria detection, using Mitotracker-647, on PN14 oocytes. In (A-C), the arrows indicate the subcortical localization, and the arrowheads depict cytoplasmic aggregation. Scale bars: 25 µm. (**D**,**E**) Electron microscopy ultrastructural analysis of WT and *Tbpl2*^-/-^ oocytes at PN7 (D) and PN14 (E). CPL: Cytoplasmic Lattices, fDy: fragmented Golgi dictyosomes, GA; Golgi Apparatus, Gm; Golgi membranes, sGm; stacked Golgi membranes, MB; Multivesicular Bodies, Mt; Mitochondria, sMt; stressed mitochondria, cRER; continuous, N; Nucleus, cRER; continuous Rough Endoplasmic Reticulum, dRER; discontinuous Rough Endoplasmic Reticulum. White arrows; mitochondria, orange arrows; RER, black arrows; Golgi membranes; yellow arrows: lattices, green arrows; multivesicular body. The asterisk (*) indicates the depletion in organelles in the subcortical region of the cytoplasm of PN14 *Tbpl2*^-/-^ mutant oocytes. The rectangles indicate the magnified field. Pictures displayed in (A-C) are single planes, scale bars: 2 µm, except in the magnification panels in (E): 1 µm.

Our electron microscopy analysis confirmed that in the small oocytes which are not yet growing, the Golgi apparatus (GA), organized in stacks of cisternae called dictyosomes, is initially located between mitochondrial clusters similarly in WT and *Tbpl2^-/-^*(Figure S3, GA). At the beginning of oocyte growth at PN7, the GA starts to reorganize in the WT cytoplasm in close proximity with mitochondria (Mt) and fragmented dictyosomes (fDy) are visible (Figure 4D, white and black arrows, respectively). Instead, in the *Tbpl2^-/-^* mutant oocytes, the cytoplasm lacks fragmented dictyosomes and is disorganized, which is further supported by the accumulation of stressed mitochondria (sMt), stacked Golgi membranes (sGM) and the appearance of multivesicular bodies (MB) in close proximity of the nucleus (Figure 4D, white, black and green arrows, respectively). Moreover, in WT oocytes, the mitochondria are reorganized by intercalation in “waves” of continuous rough endoplasmic reticulum (RER) membranes (Figure 4D, orange arrows in the magnification panels). However, no such waves are visible in the *Tbpl2^-/-^*mutant oocytes because the RER appears discontinuous and stressed mitochondria and stacked Golgi membranes (sGm) accumulate near the nucleus (Figure 4D, orange and black arrows, respectively). At PN14 in the WT growing oocytes, the cytoplasmic lattices (CPLs) are clearly formed and nicely distributed in the cytoplasm (Figure 4E, yellow arrows), as well as mitochondria and Golgi membranes (Gm) that are distributed homogeneously in the cytoplasm (Figure 4E, white and black arrows, respectively). To the contrary, in the PN14 *Tbpl2^-/-^* mutant oocytes, the cytoplasm lacks CPL and is crowded with inflated stressed mitochondria (sMt) characterized by inflated cristae, and with stacked Golgi membranes (sGm) (Figure 4E, orange and white arrows, respectively).

Altogether, these data indicate that the cytoplasmic organization is severely perturbed in the *Tbpl2^-/-^*mutant oocytes and that the cytoplasmic re-organization associated with the growing phase in WT oocytes is not occurring in the absence of TBPL2.

### Defects in the expression of key regulators associated with RNA decay

During oocyte growth, the organization of the cytoplasm is important for the storage and the regulation of the maternal transcripts. The abnormal cytoplasm organization of the *Tbpl2^-/-^* mutant oocytes strongly suggests that the regulation of RNA stability is affected. This hypothesis is supported by the transcriptome analysis of *Tbpl2^-/-^* mutant oocytes revealing that transcripts coding for key regulators of the RNA decay machinery were down-regulated (Yu *et al*, 2020). Thus, we next investigated the expression at the protein levels of several proteins associated with the RNA decay pathway. First, we analyzed the expression levels and distribution of proteins associated with the stabilization of the mRNA, such as PABPC1 which binds to the poly(A) tail in the cytoplasm (for a review, (Passmore & Coller, 2022)) and the non-specific RNA binding protein YBX2/MSY2 (hereafter called YBX2). The level of PABPC1 was increased in the mutant oocytes (Figure 5A, Figure S4A). Note that the specificity of the anti PABPC1 antibody was validated by siRNA (Figure S4B). While the levels of YBX2 localized exclusively in the cytoplasm were comparable between the WT and the mutant conditions at PN7 (Figure 5B), YBX2 aggregates were exclusively observed in the cytoplasm of mutant oocytes at PN14 (Figure 5C).

**Figure 5:**
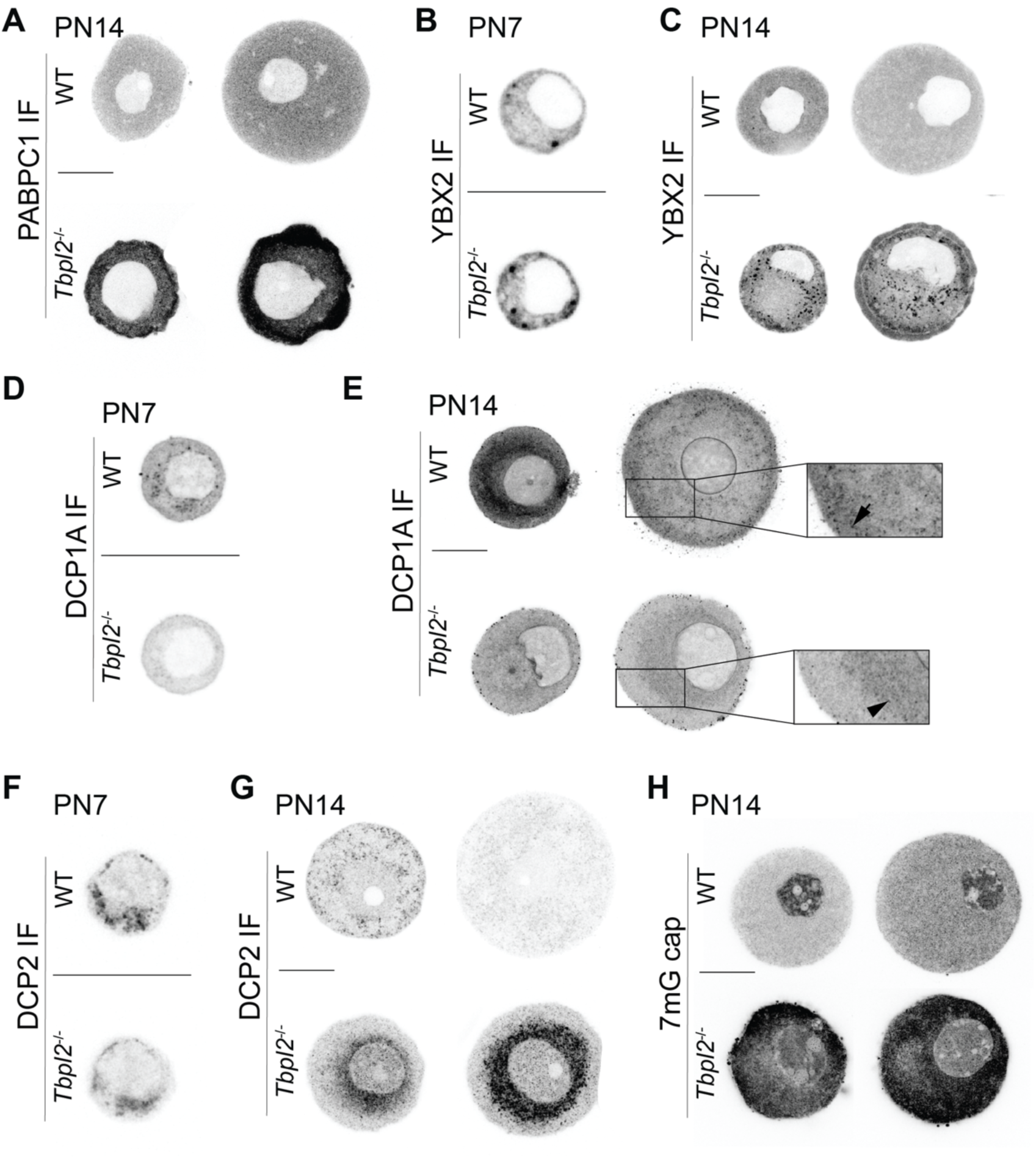
Defects in the protein complexes associated with the regulation of RNA stability. (**A-H**) Negative grayscale pictures of immunofluorescence on growing WT and *Tbpl2*^-/-^oocytes at PN7 (B,D,F) and PN14 (A,C,E,G,H) using antibodies against: PABPC1 (A), YBX2 (B,C), DCP1A (D,E), DCP2 (F,G) and 7mG cap (H). Pictures displayed are single planes, scale bar: 25 µm. In (E), the arrow indicates the subcortical accumulation in WT and the arrowhead, the nuclear juxtaposition in *Tbpl2^-/-^* mutant oocytes of DCP1A.

Second, we looked at the two subunits of the decapping complex (for a review, (Coller & Parker, 2004)); the catalytic subunit DCP2 and its mandatory partner DCP1A. Interestingly, the mRNAs coding for these proteins were thought to be dormant, meaning that they would be stored but not translated (Ma *et al*, 2013). The specificity of the anti DCP1A antibody was validated by siRNA in NIH 3T3 cells (Figure S4C). Interestingly, our data indicate that these two proteins are actually detected during oocyte growth in the control conditions (Figure 5D-G, Figure S4D). Moreover, in WT oocytes, we observed a reorganization of the distribution of DCP1A and DCP2 in the subcortical region of the oocyte at PN14 (Figure 5E,G). In the mutant oocytes, while initially no major difference could be observed at PN5 or PN7 (Figure 5D,F, Figure S4E), we could not observe the relocalization of DCP2 and DCP1A to the subcortical area at PN14 oocytes (Figure 5E,F) and surprisingly, only the expression level of DCP2 appeared highly increased in the cytoplasm in close proximity with the nucleus (Figure 5G, Figure S4F) while DCP1A levels were reduced in the cytoplasm (Figure 5E, Figure S4D), suggesting that the DCP complex may be disorganized and functionally impaired. Next, we tested whether the impairment of the decapping complex impacted the presence of capped mRNAs using an anti-7mG cap antibody in the mutant oocytes. Interestingly, we observed increased levels of 7mG cap in *Tbpl2^-/-^* mutant oocytes compared to the WT arguing for increased mRNA levels most likely owing to impaired decapping activity in the mutant oocytes (Figure 5H).

Third, we analyzed the expression of 3 subunits of the CCR4-NOT deadenylating complex: CNOT8, homolog of the yeast Caf1 deadenylase, CNOT9, homolog of the yeast Caf40 that recruits targeting partners of the deadenylase, and CNOT11 (for a review, (Caulier *et al*, 2025)). CNOT11 is a conserved subunit of the CCR4NOT complex, present in most eukaryotic organism except some fungi, that forms with CNOT1 and CNOT10 a platform for protein-protein interaction (Mauxion *et al*, 2013, 2023). Levels of these subunits were also affected in the *Tbpl2^-/-^*mutant oocytes (Figure S3G-I).

Altogether, our above data indicate that the establishment of the machinery regulating the storage and the stability of the maternal mRNAs is defective in the *Tbpl2^-/-^*mutant oocytes.

### Aberrant regulation of mRNA poly(A) tail length in the Tbpl2^-/-^ mutant oocytes

As the length of the poly(A) tail is associated with the stability of mRNAs, we carried out poly(T) FISH. As shown in Figure 6A, no difference could be detected at PN7 between WT and *Tbpl2^-/-^*mutant oocytes with a homogenous cytoplasmic signal and some foci in the nucleus. However, at PN14, we detected an increased signal in the cytoplasm of the mutant oocytes as well as in the nucleus, with the persistence and increased number as well as intensity of the nuclear foci observed earlier (Figure 6B). Quantification of the nuclear signal confirmed the increased poly(A) signal of the *Tbpl2^-/-^* mutant oocytes compared to the WT in the nucleus (Figure 6C) and in the cytoplasm (Figure S5A). As in the nucleus the poly(A) tail of the mRNAs is interacting with the PABPN1 protein, we analyzed the level of PABPN1. The distribution of PABPN1 is initially comparable between WT and mutant oocytes at PN5 and PN7 (Figure S5B). However, at PN14, we observed an increased number of bigger nuclear foci in mutant growing oocytes, (Figure 6D,E). Quantification analyses further confirmed that the increase in foci number was significant in the mutant oocytes (Figure 6E). Interestingly, the increased size of some nuclear foci, which could correspond to the PABPN1 foci, was also observed in the electron microscopy analyses (Figure 6F). We also analyzed the expression of PAPOLA, the poly(A) polymerase responsible for poly(A) tail synthesis. The expression pattern and intensity were similar at PN7 in both WT and mutant oocytes (Figure 6G), however at PN14, PAPOLA expression was more important in the *Tbpl2^-/-^* mutant oocytes (Figure 6H), both in the nucleus (Figure 6I) and in the cytoplasm (Figure S5C). Altogether, these data suggest that in *Tbpl2^-/-^* mutant oocytes, the poly(A) length of the mRNA is increased due, at least partially, to an increased activity of PAPOLA.

**Figure 6:**
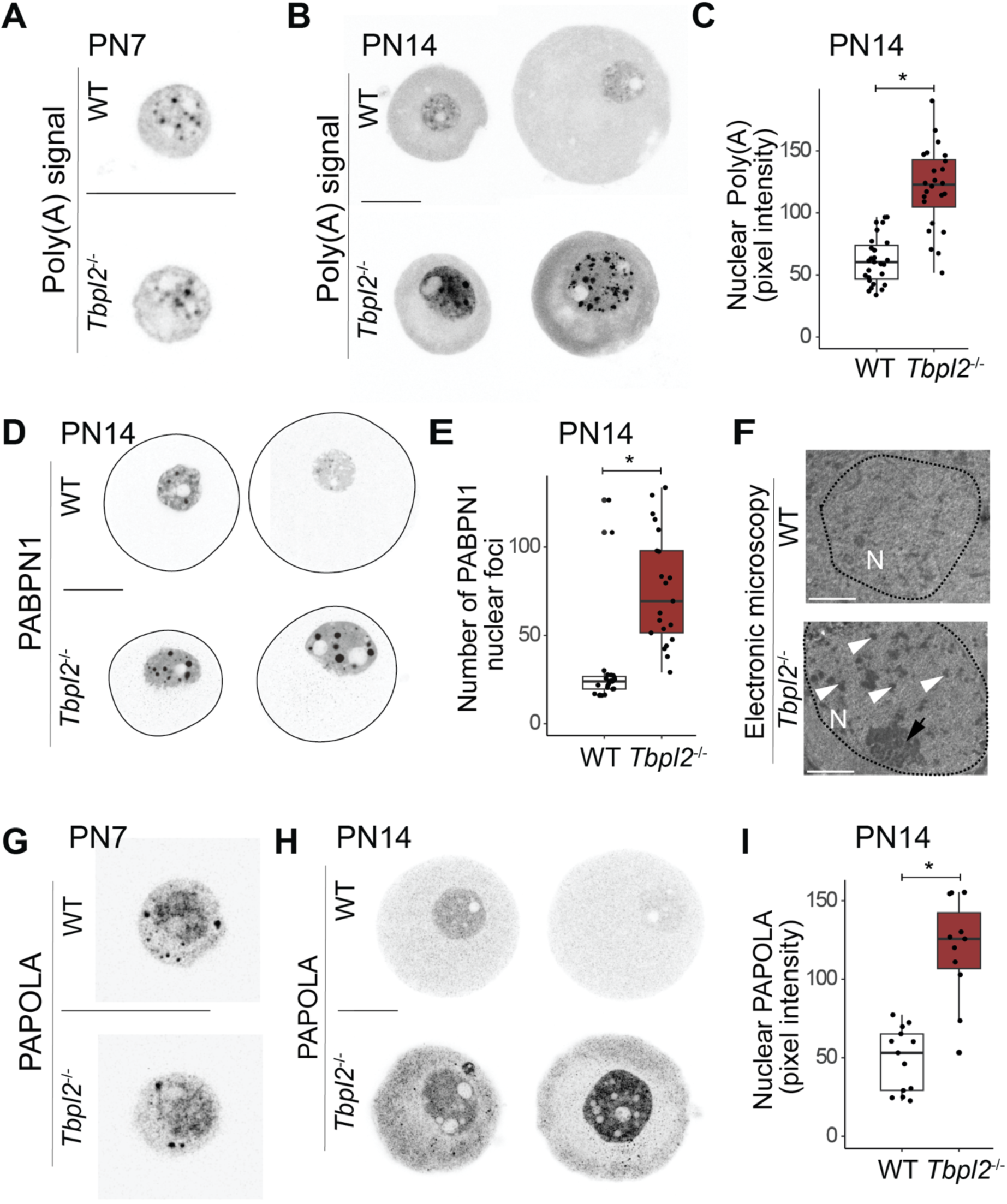
Increased poly(A) signal and expression of poly(A) associated proteins in *Tbpl2*^-/-^ mutant oocytes. (**A**,**B**) Oligo (dT) FISH on growing WT and *Tbpl2*^-/-^oocytes targeting poly(A) tails at PN7 (A) and PN14 (B). Scale bars: 25 µm. (**C**) Quantification of nuclear poly(A) signal in PN14 WT and *Tbpl2^-/-^* growing oocytes. Wilcoxon test, *: *p*-value ≤ 0.05. (**D**) Immunofluorescence on growing WT and *Tbpl2*^-/-^ mutant oocytes using an anti-PABPN1 antibody. Scale bar: 25 µm. (**E**) Quantification of the number of PABPN1 foci per PN14 oocyte nucleus. Wilcoxon test, *: *p*-value ≤ 0.05. (**F**) Ultrastructural analysis by electron microscopy of WT and *Tbpl2*^-/-^ nuclei. Scale bar: 5 µm. White arrowheads: nuclear foci, black arrow: nucleolus. (**G,H**) Detection of PAPOLA by immunofluorescence on growing WT and *Tbpl2*^-/-^oocytes at PN7 (G) and PN14 (H). Scale bars: 25 µm. (**I**) Quantification of nuclear PAPOLA in PN14 WT and *Tbpl2^-/-^*growing oocytes. The negative grayscale pictures displayed in (A,B,D,G,H) are single planes. Wilcoxon test, *: *p*-value ≤ 0.05.

### Gene specific variation of the poly(A) tail length in the Tbpl2^-/-^ mutant oocytes

To get more insight into the dysregulation of the poly(A) tail length in the absence of TBPL2, we adapted the Hairpin Adaptor-Poly(A) Tail length (HA-PAT) assay (Jiang *et al*, 2023) (Figure 7A) to measure the global distribution of the poly(A) tail. Using two sets of primers allowing to amplify the poly(A) tail length of the whole transcriptome (see Material and Methods), we confirmed that the stronger signal of poly(T) FISH observed in the *Tbpl2^-/-^*mutant oocytes correlates with an increased poly(A) smear (Figure 7B). In order to assess whether this increase is due to a lengthening of the poly(A) tail and/or a stabilization of poly(A)+ transcripts, we carried out HA-PAT on 3 selected genes, based on their regulation in absence of TBPL2 reported in (Yu *et al*, 2020). As shown in Figure 7C, we focused on *Chchd2* which is not affected, *Klf17* which is downregulated and *Uqcrc1* which is upregulated in the *Tbpl2^-/-^*mutant oocytes. These HA-PAT were performed on pools of 25-30 oocytes and in absence of reverse transcription, no signal could be detected validating the specificity of the assay (Figure 7D). HA-PAT on *Chchd2* shows comparable profile between controls and mutant samples (Figure 7E) while HA-PAT on the down-regulated transcript *Klf17* showed a clear decrease in the smear profile (Figure 7F), indicating that the poly(A) tail length of *Klf17* mRNA is decreased in absence of TBPL2. Interestingly, analysis of the up-regulated *Uqcrc1* transcript clearly showed an increased smear indicating that the poly(A) tail of *Uqcrc1* mRNA is significantly increased in the *Tbpl2^-/-^*mutant oocytes (Figure 7G), suggesting that in the absence of TBPL2, up-regulated transcripts correspond to transcripts which are stabilized by the maintenance of a long poly(A) tail.

**Figure 7:**
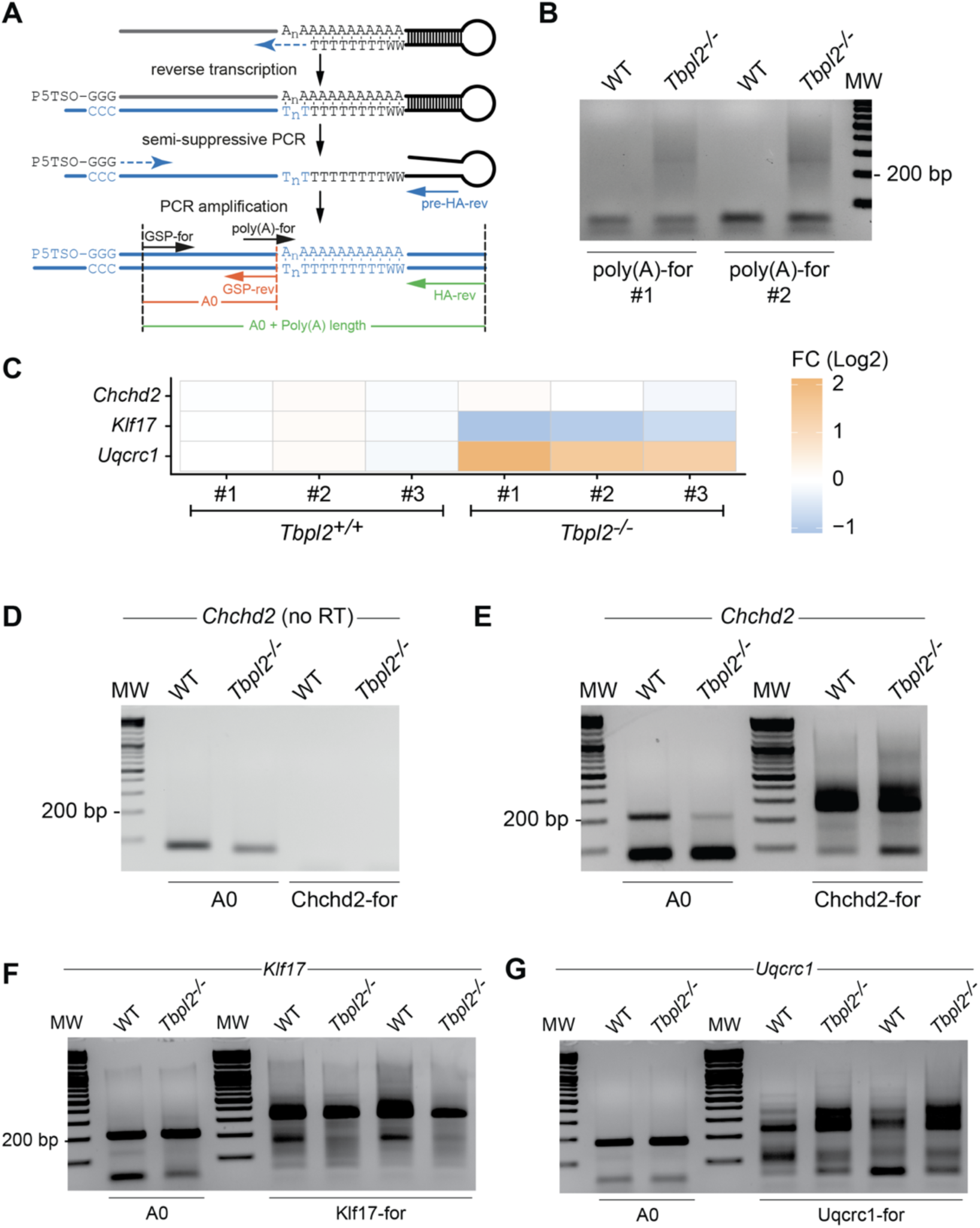
Poly(A) tails are affected in *Tbpl2^-/-^* PN14 mutant oocytes. **(A)** Schematic representation of the HA-PAT technique (adapted from (Jiang 2023)). The position of the different primers used to detect the global poly(A) tail length (poly(A)-for and HA-rev) and the gene specific poly(A) tail length (GSP-for and HA-rev) are indicated. **(B)** Evaluation of the global poly(A) tail length in 25 WT and *Tbpl2^-/-^* growing oocyte using two sets of primers indicated at the bottom of the gel. The bands below 100 bp correspond to primer dimers amplification. **(C)** Heatmap from the RNA-seq data from (Yu 2020) showing the fold change (Log2) of *Chchd2* (not significantly affected), *Klf17* (downregulated) and *Uqcrc1* (upregulated) transcripts in *Tbpl2^-/-^* mutant PN14 growing oocytes. **(D)** *Chchd2* HA-PAT control without RT. The bands correspond to the amplification of primer dimers, but there is no signal detected in absence of RT in the HA-PAT starting from 30 WT or mutant *Tbpl2^-/-^* PN14 growing oocytes. **(E-G)** HA-PAT experiment from WT or mutant *Tbpl2^-/-^* PN14 growing oocytes for the *Chchd2* (E), *Klf17* (F) and *Uqcrc1* (G) in *Tbpl2^-/-^* mutant oocytes. A0 control on the left, HA-PAT experiment on the right lanes. Thirty PN14 growing oocytes were used in (E), and 25 (left) and 30 (right) in (F,G). MW; molecular ladder, the 200 bp band is indicated.

## Discussion

In this study, we analyzed the cellular phenotype of the *Tbpl2^-/-^*mutant oocytes. In the absence of TBPL2-mediated transcription initiation, we showed that the growing oocytes fail to reorganize their cytoplasmic and nuclear compartments. The chromatin of the *Tbpl2^-/-^* mutant oocytes is less compacted and might cause the increased size of the nucleus. In the cytoplasm, the organelles and the cytoskeleton also fail to be reorganized. The disorganization of the cytoplasm is also associated with defects in the expression and localization of proteins associated with RNA stability regulation such as YBX2, involved in the stabilization of maternal RNA in oocytes (Liu *et al*, 2017). YBX2 is highly expressed in the absence of TBPL2 and could be involved in the stabilization of some transcripts. The increase in poly(A) signal observed in the absence of TBPL2-mediated transcription is also correlating with an increase in expression of the poly(A) polymerase PAPOLA and PAPBC1. Our HA-PAT analysis also suggests that in the absence of TBPL2-driven transcription in the mutant oocytes, there is an increase in the stabilization of mRNA that could be associated with an increase poly(A) length due to the reduced-functionality of the deadenylation machinery.

### Abnormal cellular organization in the Tbpl2^-/-^ mutant oocytes

In this study, we report that defects in the *Tbpl2^-/-^* mutant ovaries are detectable earlier than initially thought (Gazdag *et al*, 2009), as they are already smaller than their control counterparts at PN14. In addition, we observed that the nucleus of the *Tbpl2^-/-^* mutant oocytes are significantly bigger than the controls at all stages analyzed, from 2 weeks to 32 weeks of age. We already showed that TBPL2 is essential for the chromatin organization as shown by diminished levels of H3K4me3 and failure to reach the non-surrounded nucleolus (NSN) organization (Gazdag *et al*, 2009). Our new data confirmed that TBPL2-mediated transcription is required for compaction of the chromatin as expression of different markers associated with chromatin compaction are severely downregulated in the mutant oocytes. Interestingly, the establishment of the methylome is associated with active transcription (Veselovska *et al*, 2015). We showed that in the absence of TBPL2 mediated transcription, 5mC signal is not detected in the mutant growing oocytes. These new data strongly suggest that it is TBPL2-mediated transcription which is responsible for the establishment of the DNA methylome during oocyte growth.

In addition to the abnormal nuclear organization, we also observed alterations in cytoplasm architecture. Our analyses indicated that these defects are associated with the induction of the oocyte growth as we could not observe major phenotype at PN7 or earlier at PN5. Interestingly, this pre-growth organization seems to persist in the *Tbpl2^-/-^* mutant oocytes suggesting that the cytoplasm is not reorganized at the level of the cytoskeleton and of the – organelles. We also failed to detect the constitution of the SCMC compartment as actin failed to accumulate in the subcortical domain of the mutant oocytes, suggesting defects in regulation of actin dynamics. In our previous transcriptomic analysis, we could not detect decreased expression of *Actb*, however, expression of the positive regulator of actin polymerization, such as *Fmn2* or genes coding for the ARP2/3 complex were deregulated in the *Tbpl2^-/-^*mutant oocytes (Yu *et al*, 2020). Further analysis of the expression of these key regulators at the protein levels will be required to understand the strong decrease in cortical actin in the mutant oocytes. Altogether, our data indicate that in the absence of TBPL2-mediated transcription, the mutant oocytes are unable to engage in the growth program.

### RNA decay during oocyte growth?

The modulation of the maternal transcriptome has been well described during maturation (for reviews, see (Vastenhouw *et al*, 2019; Tora & Vincent, 2021)). As transcription stops after oocyte growth, all the subsequent events are controlled via the sequential and specific degradation of transcripts that are no longer required and/or the cytoplasmic readenylation of previously stored transcripts associated with the induction of their translation. Transcripts coding for proteins directly associated with RNA decay such as DCP1A and DCP2 (decapping complex), CNOT6L and CNOT7 (CCR4-NOT complex deadenylases) and PAN2 (PAN2/PAN3 deadenylase complex) have been categorized as dormant, i.e. stored and not translated (Ma *et al*, 2013, 2015), suggesting that the decapping activity is reduced in the growing oocytes. Due to the lack of suitable antibodies, we could not test all these proteins, but our data indicate that at least DCP1A and DCP2 are still detectable during oocyte growth, suggesting that some of the *Dcp1a* and *Dcp2* mRNA are translated or that they were translated earlier and stable. These proteins could be stored or functional during this period. We previously showed that in the *Tbpl2^-/-^* mutant growing oocytes, several transcripts coding for protein associated with the regulation of RNA stability, including *Dcp1a*, and to a lesser extent *Dcp2*, are down-regulated (Yu *et al*, 2020). Our new data showing an increase signal of 7mG cap in the mutant oocytes is in agreement with an active decapping complex activity during oocyte growth. In addition, the increase poly(A) signal we detected in the *Tbpl2^-/-^*mutant growing oocytes could also be associated with a defect in the deadenylation activity during oocyte growth. Surprisingly, while all the *Cnot* transcripts are downregulated or not changing in the *Tbpl2^-/-^*mutant oocytes (Yu *et al*, 2020), our immunolocalization analyses indicate that CNOT8, CNOT9 and CNOT11 protein levels are increased in absence of TBPL2. However, in addition to this signal increase, there is a clear change in the expression pattern, with a reduced subcortical localization and the appearance of foci, not observed in the control condition which could be associated with a defect in the deadenylation activity of the CCR4-CNOT complex. The increased poly(A) tail length we observed by HA-PAT for the upregulated *Uqcrc1* mRNA also strongly suggest that deadenylation activity is impaired in the *Tbpl2^-/-^* mutant oocytes. Interestingly, two conditional deletions of *Exosc10* coding one of the catalytic subunits of the 3’-5’ exoribonuclease exosome complex have been reported (Wu & Dean, 2020; Demini *et al*, 2023), using two different Cre lines. While Wu *et al*. used the Zp3-Cre which starts to be active at day 5, becoming fully active after the onset of oocyte growth and reported a defect in the transition between growth and maturation, Demini *et al*. used the *iGdf9-Cre* line, which is active as early as day 3 in primordial follicles oocytes and fully active at day 5 prior to the initiation of oocyte growth (Lan *et al*, 2004) and reported a role for EXOSC10 during oocyte growth. These data suggest that the contribution of the RNA decay machinery has been underestimated during oocyte growth in the mouse.

Altogether, these data strongly support that mRNA decay regulation is active and functionally important during oocyte growth and is impaired in the absence of TBPL2-mediated transcription.

### Increased poly(A) signal in the absence of TBPL2-mediated transcription

We observed an increased level of the poly(A) signal in the absence of TBPL2-mediated transcription that could be associated with the observed increase signal of the poly(A) polymerase PAPOLA and of PABPN1. We previously showed that TBP expression was decreasing during oocyte growth being replaced by TBPL2 (Gazdag *et al*, 2007) and that H3K4me3 levels and serine 2 phosphorylation of RBP1 CTD, associated with active Pol II activity were strongly reduced in *Tbpl2^-/-^* mutant growing oocytes (Gazdag *et al*, 2009), indicating that active transcription is severely impaired in the absence of TBPL2. Therefore, the increased poly(A) signal in the mutant oocytes could be the result of lack of deadenylation associated with the stabilization of an important population of mRNAs. This hypothesis is in agreement with the observation that in our previous transcriptomic analysis, 1396 transcripts were upregulated (versus 1802 transcript downregulated) (Yu *et al*, 2020) Our HA-PAT analyses interestingly suggest that there is a correlation between the impact of TBPL2 absence on mRNA levels and poly(A) tail length, further supporting our hypothesis that upregulated transcripts in *Tbpl2^-/-^* mutant oocytes are stabilized due to inefficient deadenylation. Evaluation of the impact of the lack of TBPL2-mediated transcription on poly(A) length at the level of the transcriptome by Nanopore sequencing (Lee *et al*, 2024) would be required to clarify the mechanism involved.

## Acknowledgements

We thank all members of the Vincent/Tora lab for protocols, discussions and suggestions, Dr. Maria Almonacid for critical reading and suggestions, Alicia García Sánchez for assistance with the analysis of the *Tbpl2^-/-^* sections, Sylvie Falcone, Michael Gendron and the IGBMC animal facility for the animal caretaking, the PluriCell East platform for cell culture services, the Histopathology and Embryology platform and we acknowledge the support of the Light Microscopy Facility at the IGBMC imaging center, member of the national infrastructure France-BioImaging supported by the French National Research Agency (ANR-10-INBS-04). This work was supported by funds from the Agence Nationale de la Recherche (ANR-23-CE12-0034) to SDV and Fondation pour la Recherche Médicale (EQU-2021-0312631) to SDV and LT. This work, as part of the ITI 2021-2028 program of the University of Strasbourg was also supported by IdEx Unistra (ANR-10IDEX-0002) and by SFRI-STRAT’US project (ANR 20-SFRI-0012) and EUR IMCBio (ANR-17-EURE-0023) under the framework of the French Investments for the Future Program. EGS was a recipient of a fellowship from the University of Strasbourg Doctoral School ED414.

## Author’s contributions

EGS, DD, CR, LC and CZB performed experiments. EGS and SDV designed the experiments with the help of FM and BS. SDV conceived, supervised the project and wrote the paper with support from EGS, FM, BS; DD, CR and LT.

## Material and Methods

### Animal experimentation

Animal experimentation was carried out in accordance with the ethical welfare guidelines of the French Ministry of Agriculture. The *Tbpl2* mouse line was already described (Gazdag *et al*, 2009). *Tbp2*^-/-^ male were mated with *Tbp2*^+/-^ females to generate heterozygous (considered as WT) and homozygous (KO) females.

### Histological analyses of ovaries

Ovaries were collected from post-natal day (PN) 14, and 6-, 8- and 32-weeks old WT and KO females and fixed 16 hours in Bouin’s fixative at 4°C. Ovaries were washed twice in PBS and dehydrated by immersion in successive and increasing ethanol solution concentrations (70, 90, 95 and 100%, 20 minutes each) ending in Sub-X solution (3803670, Leica Biosystems). Dehydrated ovaries were embedded in paraffin, 6 µm sections were mounted on SuperFrost Plus slides (J1800AMNZ, Epredia) and incubated 16 hours at 37°C. The sections were rehydrated in successive and decreasing ethanol solution concentrations (100, 95, 90, 70%) and stained with hematoxylin-eosin (Clinisciences K1142-1000). Image acquisition was performed with the slide scanner HAMAMATSU NanoZoomer2.0T and analysis was done using the NanoZoomer Digital Pathology view software (NDP view 3.4).

### Oocytes collection

PN5, PN7 and PN14 ovaries were dissected and adherent tissues were removed in PBS. They were incubated in a digestion buffer containing 430 µL of filtered PBS, 30 µl of collagenase (30 mg/ml, C2674-100MG, Sigma), 25 µL of hyaluronidase (10 mg/ml, H3884, type IV-S, Sigma) and 12.5 µl of trypsin (1%, 93615-5G, Sigma) The solution was incubated during 30 minutes at 37°C and homogenized each 5 minutes in order to release the oocytes. Enzymatic reaction was stopped by adding 1 ml of M2 medium (M7167-100ML, Sigma) and the oocyte suspension was transferred into a Petri dish. Oocytes were mouth-pipetted in M2 medium.

### Immunofluorescence

Oocytes were fixed 15 minutes at room temperature (RT) in a 4% PFA solution and permeabilized 20 minutes in a Blocking permeabilization Solution (BS): 0.2% Triton X-100, 1% BSA, PBS x1. Primary antibodies were diluted at 1:500 in BS and incubated overnight at 4°C in a humid chamber. Oocytes were washed twice in BS for 15 minutes each, followed by secondary antibody incubation at RT for 1 hour (dilution 1:500 in BS). Oocytes were then transferred in BS/secondary antibodies solution and incubated 1 hour at RT in a humid chamber protected from the light. Oocytes were then washed three times in BS for 10 minutes each and mounted in Vectashield-DAPI (H-1200, Vector laboratories). Antibodies references are listed in Supplemental Table 1.

### 5mC staining of oocytes

Oocytes were processed as in (Hajkova *et al*, 2010). Briefly, oocytes were fixed in 4% PFA for 20 minutes and washed three times in PBS, 1% BSA for 10 minutes. Oocytes were then permeabilized in PBS, 1% BSA, 0.5% Triton X-100 for 30 minutes, washed three times in PBS, 1%BSA for 10 minutes and treated with RNAseA (10 mg/ml) for 1hour at 37°C. Following three subsequent 10 min washes in PBS, 1% BSA, samples were rinsed in PBS and treated with 4N HCl for 20 minutes at 37°C. Oocytes were then rinsed in PBS, washed three times in PBS, 1% BSA for 10 minutes, incubated in PBS, 1% BSA, 0.1% Triton X-100 for 30 minutes and incubated in the same buffer with 5mC antibody at 4°C overnight. Oocytes were subsequently washed three times in PBS, 1% BSA, 0.1% Triton X-100 for 10 minutes and incubated with Alexa fluorophore conjugated secondary antibodies (Molecular Probes) for 1 hour at RT in the dark. Oocytes were then washed once in PBS, 1% BSA, 0.1% Triton X-100 for 10 minutes and twice in PBS, 1% BSA for 10 minutes, followed by propidium iodide (PI) staining (0.25mg/ml) for 25 minutes. The final wash was carried out in PBS, 1% BSA for 20 minutes to remove excess of PI, the oocytes were mounted in Vectashield (Vector laboratories).

### Oligo (dT) FISH

Oocytes were fixed 30 minutes at RT in a 4% PFA solution and permeabilized 20 minutes at RT in a 70% ethanol solution. Permeabilized oocytes were placed on poly-lysine coated coverslips (PB-5170, Euromedex and A38904-01, Gibco), transferred in Stellaris Wash Buffer A (SMF-WA1-60, Biosearch Technologies) supplemented with 10% formamide (F9037-100ML, Sigma) and were incubated in the dark for 16 hours at 37°C in Stellaris hybridization buffer (SMF-HB1-10, Biosearch Technologies) containing 200 nM of oligo (dT)20 cy3 probe (26-4320-02, Gene Link). Then, oocytes were washed twice at 37°C in Wash buffer A for 30 minutes each and then twice in Stellaris Wash buffer B (SMF-WB1-20, Biosearch Technologies), first at 37°C then at RT. Oocytes were mounted in Vectashield-DAPI (H-1200, Vector laboratories).

### Actin network labeling

Oocytes were fixed for 10 minutes at RT in a 4% PFA solution, and permeabilized 20 minutes in 0.2 % Triton X-100, 1 % BSA, PBS solution. Oocytes were incubated 20 minutes with 0.5 µL of Phalloidin-iFluor 488 Conjugate (23115, AAT Bioquest) in 500 µl in PBS. Then, oocytes were rinsed twice with PBS and mounted in Vectashield-DAPI (H-1200, Vector laboratories).

### Mitochondria labelling

Oocytes were incubated 20 minutes with 500 nM of Mitotracker Deep Red FM (8778, Cell Signaling) in M2 medium (M7167-100ML, Sigma). Oocytes were then fixed for 30 minutes at RT in a 4% PFA solution, rinsed twice with PBS and mounted in Vectashield-DAPI (H-1200, Vector laboratories).

### Imaging and fluorescence quantifications

Samples were imaged using Leica inverted confocal microscope with a HC PL APO CS2 63x/1.4 oil and 10x objective. Mean nuclear fluorescence intensity was quantified using ImageJ (version 1.54t). For each nucleus, fluorescence intensity was measured in several regions of interest (ROIs) of known size. The mean fluorescence intensity values obtained from the different ROIs were averaged to obtain the mean nuclear fluorescence intensity for each nucleus. Nuclear signal was normalized on the cytoplasmic signal. Statistical analysis and plot were generated using R (version 4.5.2) and the ggplot2 package (version 4.0.3). β-TUBULIN signal was quantified by extracting the entire cytoplasmic intensity of each individual oocyte. Subcortical ACTIN signal was quantified by extracting the signal form the cytoplasmic periphery of each oocyte.

### Electron microscopy

The samples were fixed by immersion in 2.5% glutaraldehyde and 2.5% paraformaldehyde in cacodylate buffer (0.1 M, pH 7.4) and washed in cacodylate buffer for further 30 minutes. Oocytes were post fixed in 1% osmium tetroxide in 0.1 M cacodylate buffer for 1 hour at 4°C and dehydrated through increasing ethanol solution concentrations (50, 70, 90, and 100%) and propylene oxide for 30 minutes each. Samples were embedded in Epon 812 (Euromedex EM-14900), and 70 nm ultrathin sections were generated (Leica Ultracut UCT), contrasted with uranyl acetate and lead citrate and examined at 70kv with a Morgagni 268D electron microscope (FEI Electron Optics, Eindhoven, the Netherlands). Images were captured with Mega View III camera (Soft Imaging System).

### NIH3T3 culture and siRNA validation

NIH3T3 cells were cultured on coverslips in DMEM (10% NCS supplemented) for 70% confluency and conditioned 2 hours in Opti-MEM reduced serum medium (31985062, Gibco). ON-TARGETplus (Dharmacon) siRNA were diluted with Lipofectamine (52887, Invitrogen) in Opti-MEM to 50 nM. NIH3T3 cells were silenced during 48h, fixed with 4 % PFA for 15 minutes, washed twice with PBS and permeabilized with 0.5% Triton X-100 for 20 minutes. Antibodies were diluted 1:500 in Blocking permeabilization Solution (BS, 0.2 % Triton X-100, 1 % BSA, PBS x1) and incubated overnight at 4°C. Cells were washed twice with PBS and secondary antibodies were incubated 1 hour at RT. Coverslips were washed twice and mounted in Vectashield-DAPI (H-1200, Vector laboratories).

### Hair-pin Adaptor Poly(A) Tail length (HA-PAT)

The protocol was carried out as described in the original publication (Jiang *et al*, 2023) for the gene specific HA-PAT and adapted for the global poly(A) tail length. Briefly, pools of 25-30 PN14 oocytes were collected in M2 medium, washed, and transferred to 2 µl lysis buffer (0.2% Triton X 100, 2 U/µl RNase inhibitor) in 0.2 ml PCR tubes, incubated on ice for 10 min, and stored at -80 °C until use. Lysates (2 µl) were denatured at 72 °C for 3 min and placed on ice, then incubated with 1 µl of annealed hairpin adaptor (10 µM) at 25 °C for 10 min to allow hybridization of the hairpin to the poly(A) tails; all primer and hairpin adaptor sequences are listed in Supplemental Table 2. Reverse transcription was carried out at 42 °C for 90 min in a 20 µl reaction containing SuperScript IV buffer (18090010, Invitrogen), 5 mM DTT, 60 mM MgCl_2_, 1 M betaine (B0300, Sigma), 1 mM dCTP (18253013, Fisher Scientific), 0.5 mM dNTPs (04728858001, Roche), 1.5 µM P5TSO primer, 10 U RNAsin Ribonuclease Inhibitor (N2511, Promega), and SuperScript IV reverse transcriptase. Semi-suppressive PCR was then performed using P5TSO and Pre-HA-rev primers with AmpliTaq Gold 360 (4398881, Applied Biosystems), in a 50 µl reaction and a cycling program of 95 °C for 1 min; 16 cycles of 95 °C for 15 s, 65 °C for 10 s, and 68 °C for 2 min. Preamplified cDNA was stored at -20 °C. Gene-specific PCRs were carried out using either gene-specific forward and reverse primers (A0 amplicon) or a gene-specific forward primer and HA-rev primer (poly(A) amplicon), with 2 µl of preamplified cDNA, a standard PCR mix, and cycling conditions of 94 °C for 2 min; 35 cycles of 94 °C for 30 s, 62 °C for 30 s, and 72 °C for 30 s; followed by a 7-min extension at 72 °C. PCR products (10 µl) were resolved on 2% agarose gels in 1x TBE at 100 V for 35 min to assess poly(A) tail length distributions.

## Supplemental Figures

**Supplemental Figure 1:**
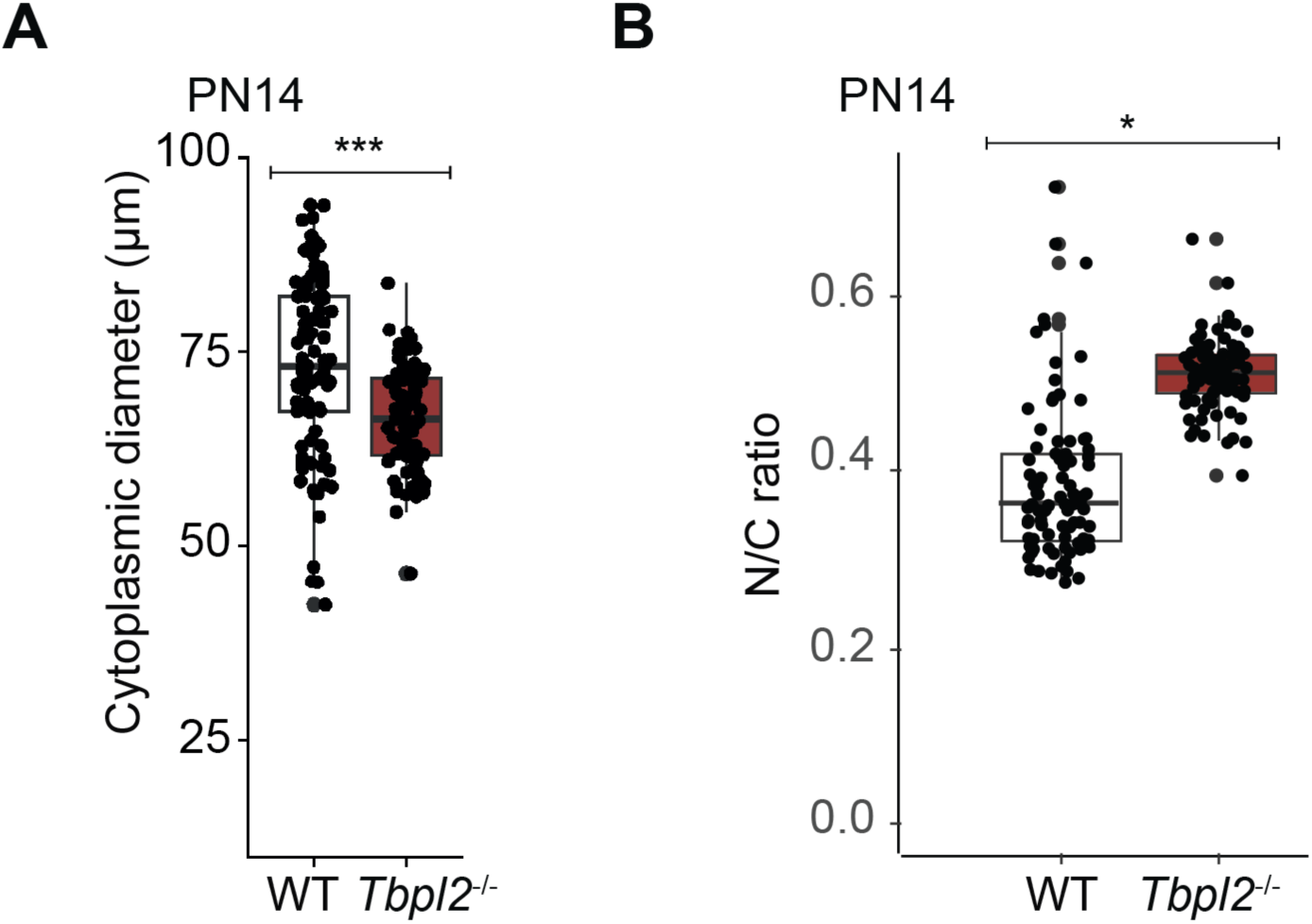
The *Tbpl2*^-/-^ mutant growing oocytes are smaller but have a bigger nucleus. (**A**) Quantification of the cytoplasmic diameter of PN14 isolated growing oocytes. (B) Quantification of the nuclear to cytoplasm (N/C) ratio of PN14 isolated growing oocytes. Wilcoxon test, *: *p*-value ≤ 0.05, **: *p*-value ≤ 0.01, ***: *p*-value ≤ 0.001.

**Supplemental Figure 2:**
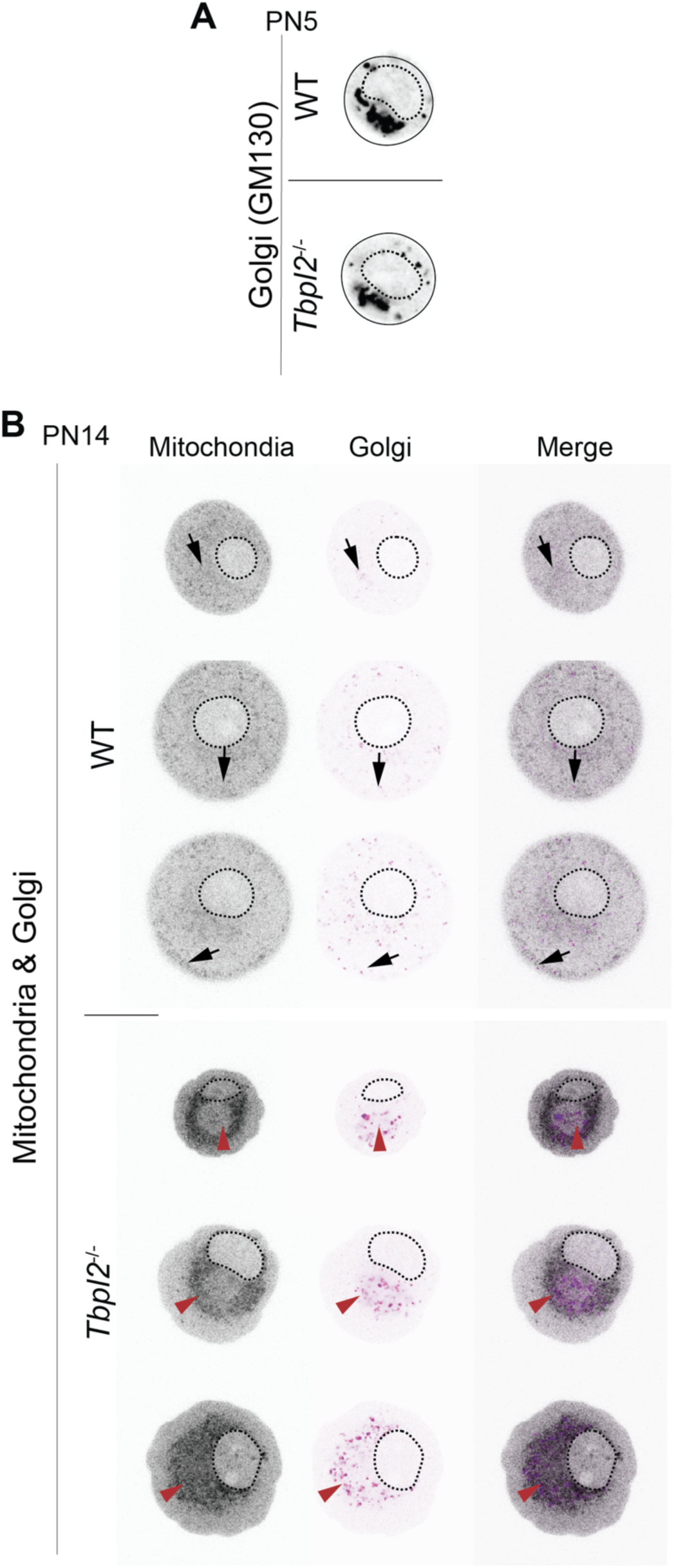
Subcellular localization of Golgi and mitochondria. (**A**) Immunofluorescence on WT and *Tbpl2*^-/-^ mutant oocytes using antibody against the Golgi protein GM130 at PN5. (**B**) Codetection of the Golgi (anti-GM130 immunostaining) and of the mitochondria using Mitotracker-647 at PN14. Black arrows: cytoplasmic distribution of Golgi and mitochondria, red arrowheads: cluster of Golgi and mitochondria in close apposition to the nucleus in the mutant *Tbpl2^-/-^* oocytes. Scale bar: 25 µm.

**Supplemental Figure 3:**
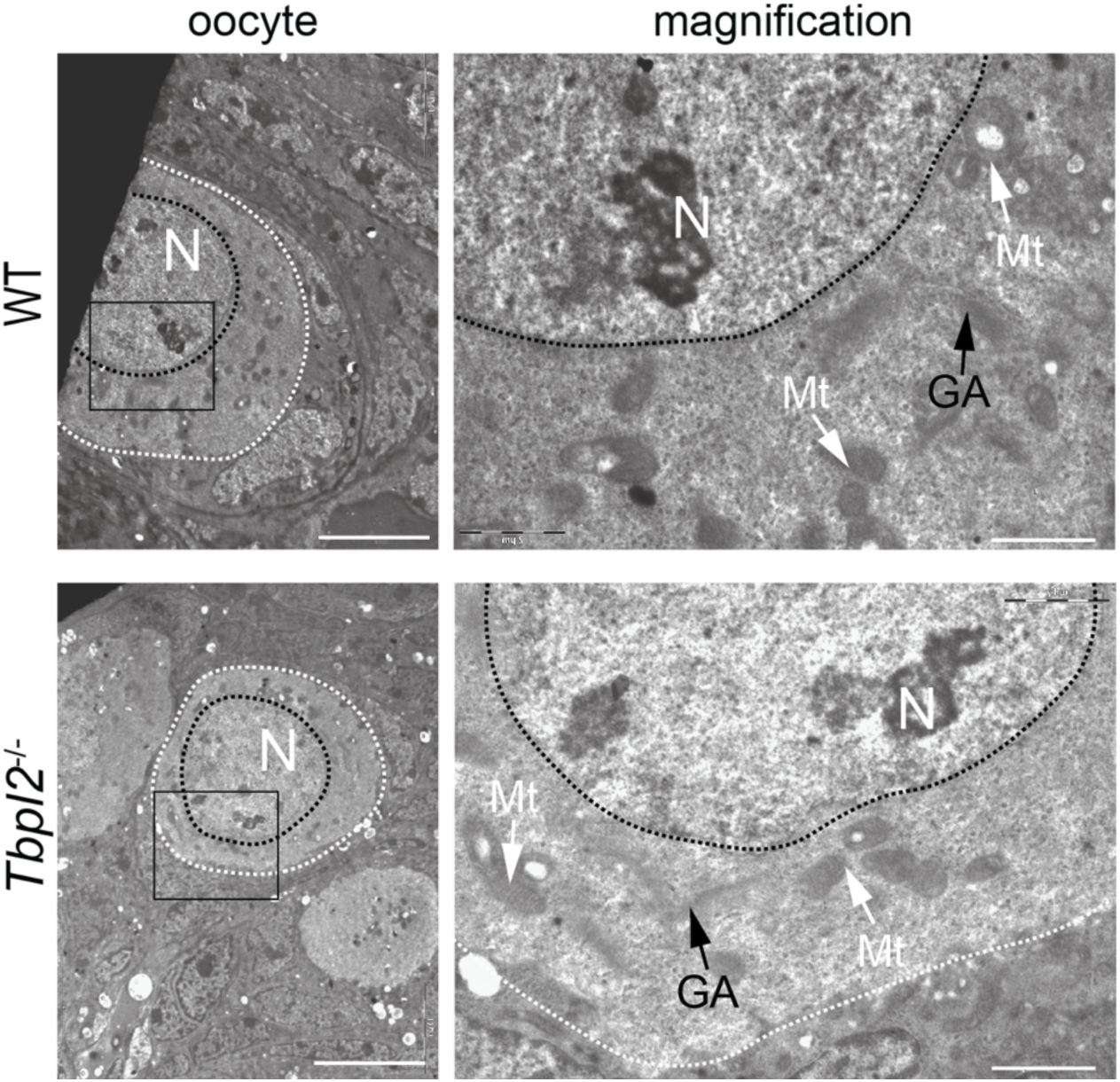
Additional ultrastructural analysis of the subcellular distribution of organelles. Ultrastructural analysis by electronic microscopy of the subcellular organization of a small oocyte not yet growing at PN14. Magnified fields are indicated by rectangles. White and black dashed lines, respectively highlight the oocyte and nucleus delimitations. N: Nucleus, GA; Golgi Apparatus, L: Lattices, Mt: Mitochondria. White arrows; mitochondria, black arrows; Golgi membranes. Scale bars: 10 µm, 2 µm for magnifications.

**Supplemental Figure 4:**
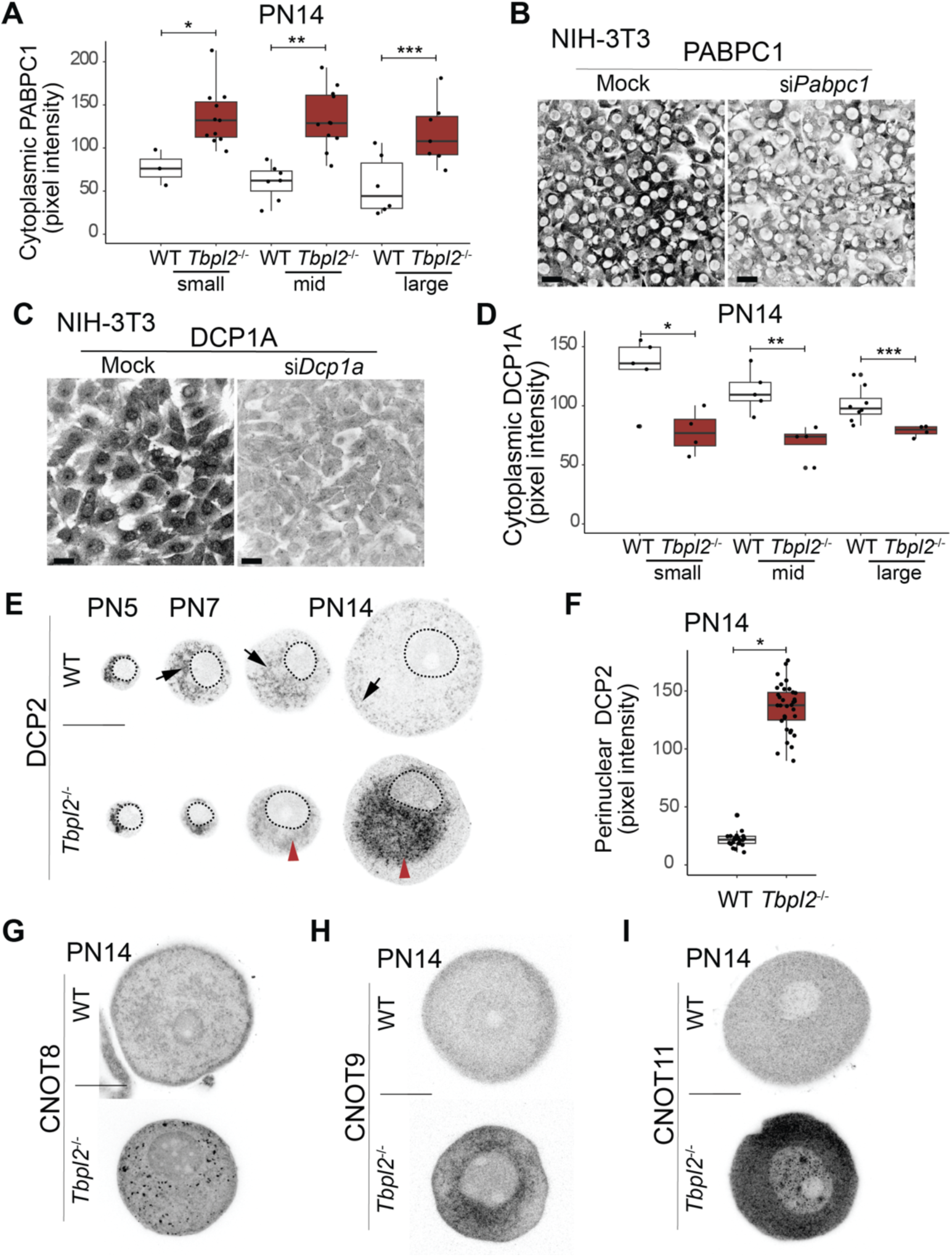
Analysis of the expression of subunits of protein complexes associated with the regulation of RNA stability. (**A**) Quantification of cytoplasmic immunostaining of PABPC1 in PN14 WT and *Tbpl2^-/-^* growing oocytes according to their size (small, mid and large). (**B,C**) Immunofluorescence detection of DCP1A (B) and PABPC1 (C) on 3T3 cell line after siRNA treatment (mock and si*Dcp1a* (B) or *siPabpc1* (C)). Scale bars: 15 µm. Scale bars: 15 µm. (**D**) Quantification of cytoplasmic immunostaining of DCP1A in PN14 WT and *Tbpl2^-/-^* mutant oocytes. (**E**) Analysis of DCP2 expression at different developmental time points (PN5, PN7 and PN14) by immunofluorescence in WT and *Tbpl2^-/-^*mutant oocytes. The big WT oocytes are underrepresented in the ovary at PN7 but are depicted in order to show that the *Tbpl2^-/-^* phenotype is first detected at that stage. (**F**) Quantification of DCP2 perinuclear immunostaining in PN14 WT and *Tbpl2^-/-^* mutant oocytes. (**G**-**I**) Immunofluorescence on PN14 growing WT and *Tbpl2*^-/-^ mutant oocytes using antibodies against CCR4-NOT subunits CNOT8 (G), CNOT9 (H) and CNOT11 (I). Scale bars: 25 µm. Wilcoxon test, *: *p*-value ≤ 0.05, **: *p*-value ≤ 0.01, ***: *p*-value ≤ 0.001.

**Supplemental Figure 5:**
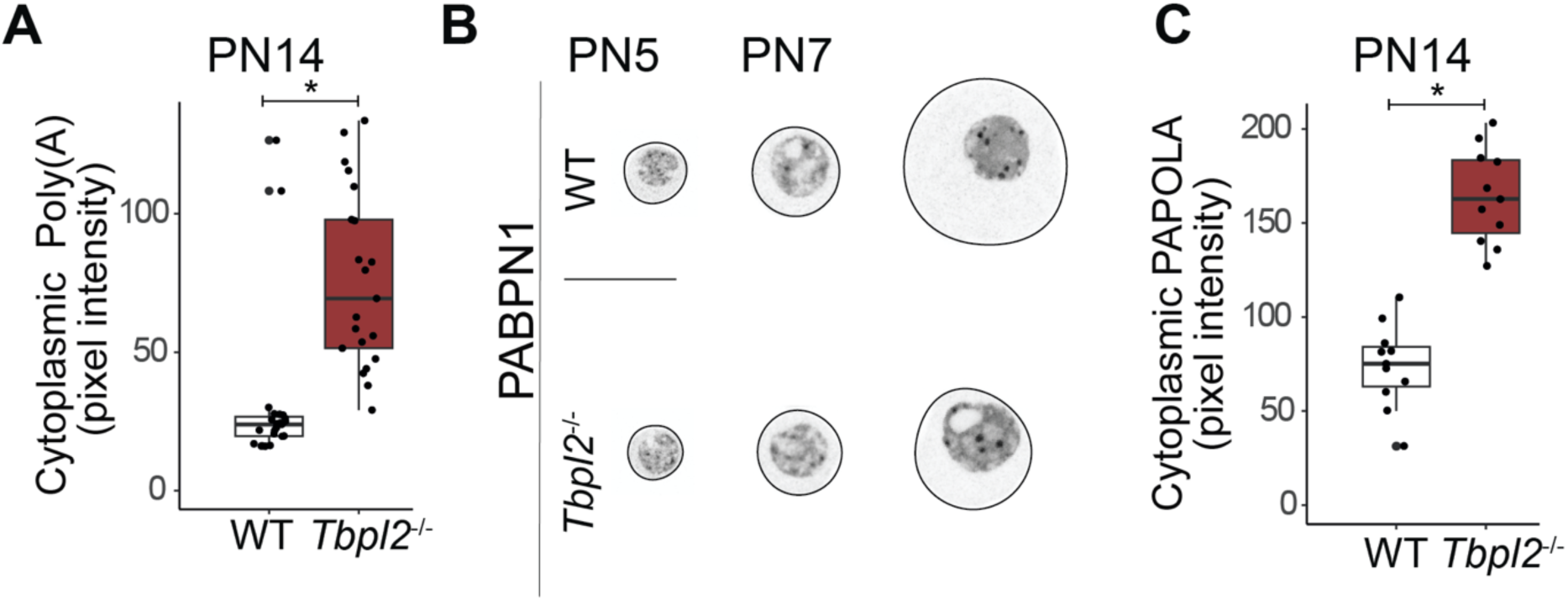
Additional quantifications and analysis of poly(A) tail and poly(A) associated proteins. (**A**) Quantification of cytoplasmic poly(A) signal in PN14 WT and *Tbpl2^-/-^* mutant oocytes. (**B**) Analysis of PABPN1 expression at two developmental PN5 and PN7 by immunofluorescence in WT and *Tbpl2^-/-^* mutant oocytes. The big WT oocytes are underrepresented in the ovary at PN7 but are depicted in order to show that the *Tbpl2^-/-^* phenotype is first detected at that stage. (**C**) Quantification of cytoplasmic immunostaining of PAPOLA in PN14 WT and *Tbpl2^-/-^* growing oocytes. Wilcoxon test, *: *p*-value ≤ 0.05.

**Supplemental Table 1:**

| Antibodies | Source | reference |
| --- | --- | --- |
| Mouse monoclonal anti-5mC | Diagenode | C15200081-100 |
| Mouse monoclonal anti-7mG | Sigma | MABE419 |
| Mouse monoclonal anti-β-TUBULIN | IGBMC | 1TU-2A2 |
| Mouse monoclonal anti-GM130 | BD BioSciences | 610822 35/RUO |
| Mouse monoclonal anti-CNOT8 | T. Yamamoto, OIST,<br>Japan | (Takahashi *et al*, 2015) |
| Rabbit polyclonal anti-CNOT9 | H. Okayama, Tokyo<br>University, Japan | αRcd-C, (Hiroi *et al*,<br>2002) |
| Rabbit polyclonal anti-CNOT11 | IGBMC | 3025, (Mauxion *et al*,<br>2013) |
| Rabbit polyclonal anti-DCP1A | IGBMC | 90, (Dijk *et al*, 2002) |
| Rabbit polyclonal anti-DCP2 | Invitrogen | PA5-115102 |
| Rabbit polyclonal anti- DESMIN | Abcam | ab15200 |
| Mouse monoclonal anti-GM130 | BD BioSciences | 610822 35/RUO |
| Rabbit polyclonal anti-H3K27me3 | Millipore | 07-449 |
| Rabbit polyclonal anti-H3K9me3 | Diagenode | C15200146-10 |
| Rabbit polyclonal anti-HP1β | Cell signaling | 8676T |
| Rabbit polyclonal anti-PABPC1 | Abcam | ab21060 |
| Rabbit polyclonal anti-PABPN1 | Bethyl laboratories | A303-523A |
| Rabbit polyclonal anti-PAPOLA | Abcam | ab251660 |
| Rabbit polyclonal anti-YBX2 | Abcam | ab33164 |
| Goat anti-mouse; Secondary Antibody 488 | Thermo Scientific | A32723 |
| Goat anti-rabbit; Secondary Antibody 594 | Thermo Scientific | A-11037 |

**Supplemental Table 2:**
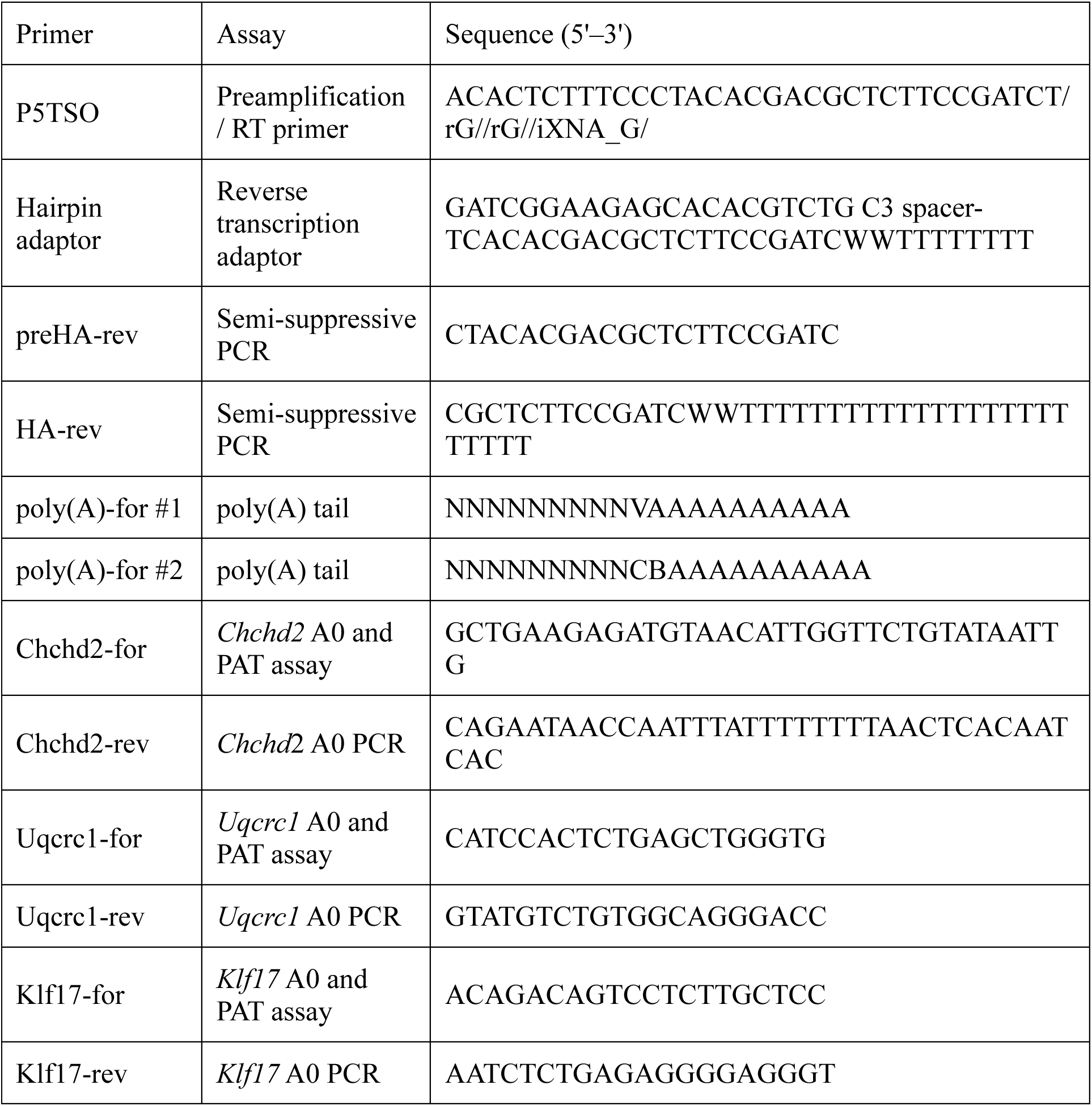

